# Neural Encoding of Semantic Structures During Sentence Production

**DOI:** 10.1101/2024.11.22.624906

**Authors:** Laura Giglio, Peter Hagoort, Markus Ostarek

**Affiliations:** Max Planck Institute for Psycholinguistics, Netherlands; Donders Institute for Brain, Cognition and Behaviour, Netherlands; Department of Communication Sciences and Disorders, Arnold School of Public Health, University of South Carolina, Columbia SC, USA

**Keywords:** compositional processing, fMRI, sentence production, language, thematic roles

## Abstract

The neural representations for compositional processing have so far been mostly studied during sentence comprehension. In an fMRI study of sentence production, we investigated the brain representations for compositional processing during speaking. We used a rapid serial visual presentation sentence recall paradigm to elicit sentence production from the conceptual memory of an event. With voxel-wise encoding models, we probed the specificity of the compositional structure built during the production of each sentence, comparing an unstructured model of word meaning without relational information with a model that encodes abstract thematic relations and a model encoding event-specific relational structure. Whole-brain analyses revealed that sentence meaning at different levels of specificity was encoded in a large left fronto-parieto-temporal network. A comparison with semantic structures composed during the comprehension of the same sentences showed similarly distributed brain activity patterns. An ROI analysis over left fronto-temporal language parcels showed that event-specific relational structure above word-specific information was encoded in the left inferior frontal gyrus (LIFG). Overall, we found evidence for the encoding of semantic meaning during sentence production in a distributed brain network and for the encoding event-specific semantic structures in the LIFG.

## Introduction

One of the most characteristic features of human language is its almost infinite combinatorial potential. The same words can be used in different structures and combinations to compose different meaning. For example, the sentences “the surfer sees the violinist” and “the violinist sees the surfer” are formed by the same words, but have different meaning that is determined by the relational structure of the event. Similarly, in “the surfer sees the violinist” and “the surfer is seen by the violinist”, the content words are the same and in the same order, but the voice of the sentence (active vs. passive structure) indicates a different relational structure and meaning. Sentence-level meaning is thus composed by a combination of word meanings which forms a semantic structure based on syntactic rules to characterize a conceptual event representation (Jackendoff 1992). Investigations on how sentence-level compositional meaning is supported by brain structure and function have only just started.

At the phrasal level, many MEG studies found that the left anterior temporal lobe (LATL) and the ventro-medial prefrontal cortex (mPFC) sequentially activate for semantic composition (e.g. composing “red boat” vs. “red blue”) (e.g. Bemis & Pylkkänen, 2011; Blanco-Elorrieta et al., 2018; Flick et al., 2018; Pylkkänen, 2020; Pylkkänen et al., 2014; but see Kochari et al., 2021). At the sentence level, the argument structure of the verb defines the thematic roles of the entities participating in an event (e.g. *who did what to whom*). The angular gyrus, extending to posterior superior temporal gyrus (STG) and middle temporal gyrus (MTG), shows increased activation for verbs with more complex verb argument structures (Meltzer-Asscher et al. 2013; Meltzer-Asscher et al. 2015), and is involved in event processing (Matchin et al. 2019). In addition, the mid left MTG was found to decode the thematic roles of nouns in a video depicting an action event, where a rabbit “punching a monkey” was classified as an agent vs. a patient (Wang et al. 2016). An adjacent region in mid left STG was instead found to decode the identity of agents and patients (i.e. determining if the agent was a rabbit or a monkey) (Frankland and Greene 2015; Frankland and Greene 2020a). Lesions to these mid regions of temporal cortex were also found to lead to deficits in accessing thematic role knowledge at the linguistic and conceptual level (Wu et al. 2007). Therefore, the mid posterior temporal lobe, extending to the angular gyrus, is consistently linked with processing of thematic roles and verb argument structure.

There is an advantage for modelling compositional effects at the sentence level, relative to individual word meanings. Constraining the meaning of a word in a sentence by verb semantics provides a better fit to brain data relative to its meaning out of context, and this incremental composition is seen to occur in a parieto-temporo-frontal network (Lyu et al. 2019). Sentence-specific propositional meaning best models brain activity relative to unstructured models that use individual word meanings separately (bag-of-words) and is found to be distributed across the brain and language network (Anderson et al. 2021). Regarding *how* compositional meaning may be supported by brain structure, Frankland & Greene (2015) suggested a neural architecture where regions of the cortex flexibly encode the values of semantic variables (such as agent and patient). In particular, they found that a region of the left mid superior temporal cortex (lmSTC) can distinguish between mirror sentences, which use the same words to create different meaning (e.g. “the surfer sees the violinist” and “the violinist sees the surfer”) and carries information about the identity of the actors of an event to the amygdala. Additionally, adjacent subregions of the lmSTC separately carry information about the identity of the agent and patient of events, showing that agent and patient roles can be encoded simultaneously as abstract semantic variables to potentially form complex semantic representations. Moreover, the anterior medial prefrontal cortex (amPFC) and the hippocampus encode verb-specific information about the thematic roles of entities participating in an event, i.e. thematic roles in combination with a specific event (such as surfer-as-seer, Frankland and Greene 2020a). The results of these studies were integrated to propose a functional architecture that dynamically encodes sentence meaning in the brain relying on neural encoding in the amPFC, lmSTC and hippocampal (Frankland and Greene 2020b).

In the current study, we asked how compositional sentence meaning is encoded during sentence production in the brain. Semantic structures are thought to be shared between production and comprehension (Levelt 1989; Guhe 2007). However, sentence production and comprehension have different timing demands (Momma and Phillips 2018; Giglio, Ostarek, et al. 2024). In comprehension, the compositional meaning represents the end goal, while in production it is the prerequisite, but its unfolding dynamics are unclear. A conceptualizer system generates a preverbal (or prelinguistic) message in the form of a structured conceptual representation (Levelt 1989; Jackendoff 1992; Guhe 2007). The conceptual structure is then formulated into a linguistic linear sequence that eventually takes the form of a phonetic plan for the articulators (Levelt 1989). The formulation of conceptual structures into linguistic sequences requires semantic concepts (entities and events), as well as the semantic roles between concepts (the argument structure) and features (e.g. number) to be encoded during message generation (Guhe, 2007). The semantic roles that take part in the event are mapped to lexical items assigned to syntactic roles (e.g. subject, object, verb; for a non-lexicalist account, see Krauska and Lau 2023). The syntactic roles are then ordered into a phrase-structure frame that includes morphological and phonological specification (Bock and Levelt 1994).

One critical characteristic of sentence production is its incrementality at different levels (e.g. Bock & Ferreira, 2014; Levelt, 1989). Parts of the message can thus be formulated (e.g. lexically encoded) before the whole message is generated (De Smedt and Kempen 1987). Similarly, formulation proceeds incrementally and can continue while words are being articulated. Therefore, the composition of complex semantic structures may proceed while sentence production unfolds, instead of being fully defined before the start of the sentence. The accessibility of lexical items and the codability of the events to be described have been found to affect the order in which entities are uttered during sentence production (Kuchinsky et al. 2011; van de Velde et al. 2014). Similarly, the verb is not always planned before sentence onset (Momma et al. 2016; Momma and Ferreira 2019). As a consequence, compositional representations at different levels of abstraction may be built at different time points between production and comprehension (see for different dynamics during syntactic processing in production and comprehension, Giglio, Ostarek, et al. 2024).

Despite these timing differences between production and comprehension, picture description and eye-tracking studies have highlighted that there is an initial point where the message is globally encoded during production. For example, studies taking advantage of picture descriptions have found that message generation is affected by the complexity of the event. In particular, evidence on speech onset times shows that longer utterances have slower onset latencies. For instance, for the sentence “The dog and the foot move up and the kite moves down” the onset is later than “the dog and the kite move up”, even though the subjects are equally long, suggesting that there is high-level processing of an event for the remainder of the sentence before speech onset (Smith and Wheeldon 1999). Eye-tracking studies show that there is an initial central fixation of 300-400ms for higher-level coding of the scene during picture description experiments before attention is moved to the first item to be uttered (Griffin and Bock 2000; Konopka 2019). Konopka (2019) showed that the event is encoded before speech onset by finding that speech onset fixations are directed to the character that is most informative for action encoding, providing evidence for early event encoding during conceptualization. In addition, during masked scene processing participants can rapidly identify the agents and patients of the depicted action as measured by questions following the rapid scene presentation (Hafri et al. 2013). Therefore, entities participating in an event seem to be connected to their thematic role in very short processing timescales during scene processing (cf. Grimshaw, 1990; Jackendoff, 1992). Overall, evidence from scene description experiments suggests that the event is encoded early and that eye movements roughly follow identities in a scene in the order of formulation and articulation (see for a review Papafragou & Grigoroglou, 2019).

Therefore, psycholinguistic evidence based on scene description experiments suggests that compositional structures are built early and quickly in production. Advances in neuroimaging, especially decoding and feature-based encoding models, have made it possible to probe neuro-cognitive processes that unfold during language planning and production. Frankland and Greene (2020a) found evidence for both abstract representations (across event types) and specific representations (event/verb-specific) to be encoded in the brain, in different regions, during sentence comprehension. Here, we asked whether it would be possible to probe brain activity patterns for semantic structures composed during sentence production, and, if so, whether they would be encoded in separate brain regions as found by Frankland and Greene (2020a) or in distributed brain activity as suggested by several other studies (Blank et al. 2016; Anderson et al. 2019; Lyu et al. 2019; Anderson et al. 2021).

Previous studies found that a similar network is engaged during sentence production and comprehension with some asymmetries in the level of engagement. In particular, both temporal and frontal areas are involved in sentence production (Segaert et al. 2012; Matchin and Wood 2020; Giglio et al. 2022; Hu et al. 2022; Giglio, Sharoh, et al. 2024), although production and comprehension differ in how much they activate them (Giglio et al. 2022; Arvidsson et al. 2024). These results thus indicate that the neural resources of production and comprehension are likely shared, and as such provide the preconditions to expect the same or a similar network for compositional processing in production and comprehension. Therefore, in the current study, we investigated the network encoding compositional structure at different levels of abstraction. We investigated the specificity of relational encoding using voxel-wise encoding models trained to predict brain activity from sentence descriptors positing different levels of specificity in noun-role combinations. We ran whole-brain analyses to determine which regions were involved in compositional processing in production. We then explored how brain parcels for an extended language network (e.g. Hu et al. 2022) encoded different aspects of sentence meaning to isolate regions encoding event-specific compositional meaning over and above event-general thematic roles.

## Materials and Methods

### Participants

Forty right-handed native Dutch speakers participated in the study after giving written informed consent (27 females, mean age = 21.5 years, range 18-49). The study was approved by the ethical committee for Region Arnhem Nijmegen. Participants reported having no language-related or neurological disorders and normal or corrected-to-normal vision. Two participants were excluded due to poor performance during the task (n = 1 not enough trials per run for the analysis; n = 1 question answering at chance).

### Materials

The stimuli consisted of 252 Dutch sentences made of a combination of 16 nouns and 16 verbs; an additional 36 sentences of the same type followed by a question; and 72 filler sentences. Half of the sentences of interest were in active voice, while the other half was in passive voice, half of which in one word order and half in another word order (allowed in Dutch: “de bokser wordt door de muzikant herkend”, “the boxer is by the musician recognized” or “de bokser wordt herkend door de muzikant”, “the boxer is recognized by the musician”). The filler sentences were composed of several verbs and nouns to increase variability of content and structure in the production. The use of sentences in both active and passive voice allowed us to focus on the encoding of thematic roles independently from the position in the sentence (as the order of thematic roles is swapped in active and passive sentences).

To probe the encoding of different roles in the sentence, we used nouns that referred to individuals playing either sport or music, 8 for each category (see Table 1). The verbs were all transitive verbs either expressing an action that involves contact, or that requires perception. All perception verbs were experiencer-subject verbs, with the perceiver (i.e. experiencer) expressed in subject position in active sentences and the perceived item (i.e. stimulus) as a grammatical object (e.g. “the athlete sees the musician”). The experiencer-subject term refers to verbs used in active voice: in passive sentences the perceived item (stimulus) is expressed as subject. The two categories of nouns and verbs were matched in length and frequency (see Table 1) (Keuleers et al. 2010). We contrasted semantic categories rather than individual verbs and nouns to increase variability in the production output and to avoid too much word repetition. Both noun and verb categories were found to be decodable before (Xu et al. 2018; Arana 2022). All possible noun-verb-noun combinations led to a pool of 3840 sentences (with two nouns in each sentence, excluding sentences with identical subjects and objects). We thus made a selection of sentences to ensure that all verbs were used the same number of times, and nouns occurred the same number of times in agent and patient roles. We additionally ensured that each noun was combined with each verb at least once in each role, and kept as few subject-object identical noun combinations as possible. Finally, there were equal numbers of sentences for the combination of each noun semantic category and each verb semantic category (eight combinations, e.g. musician-agent, verb-contact, musician-patient: 36 sentences). Sentences could have two nouns of the same category and of different categories (but never identical nouns). The list of sentences used can be found on the Radboud Data Repository.

**Table 1:**
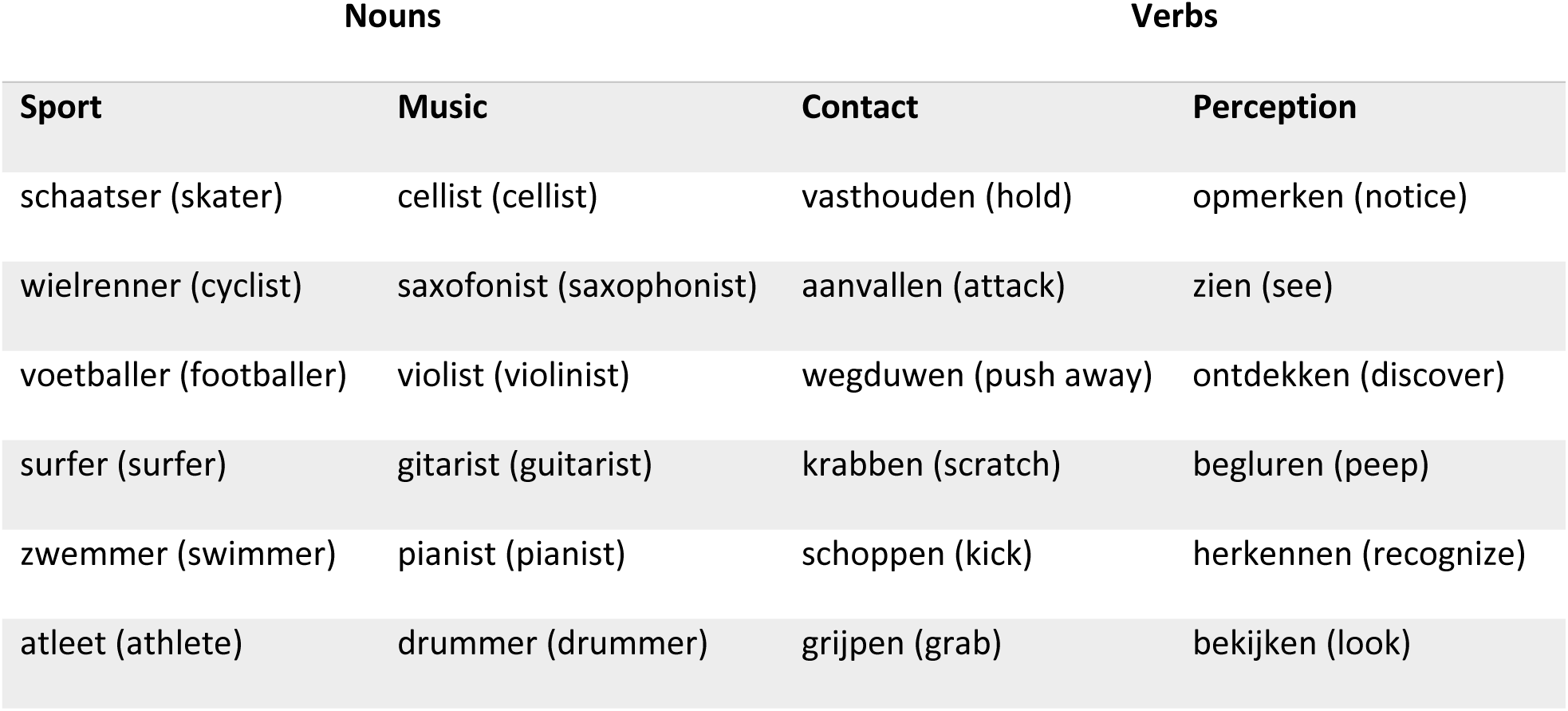
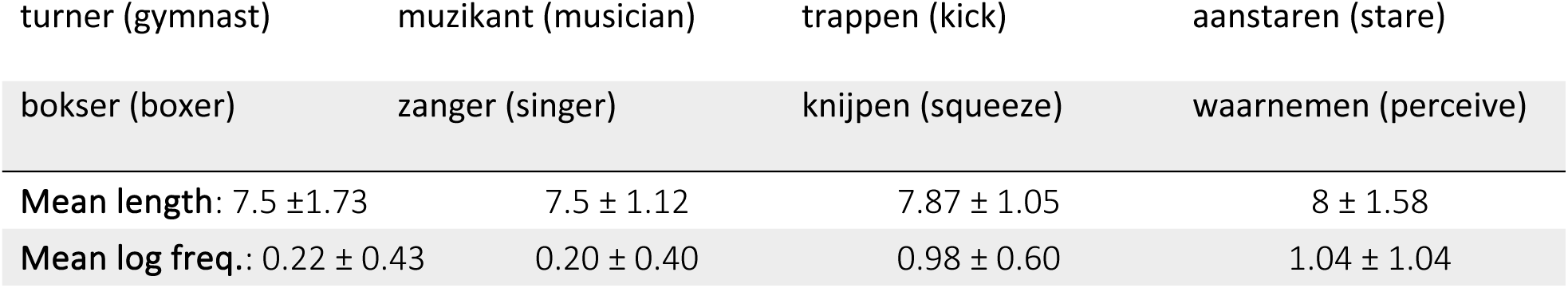
Nouns and verbs used in different combinations to create the sentences of interest, with their mean length and log frequency values ± standard deviation.

### Procedure

The experiment used a rapid serial visual presentation (RSVP) sentence recall paradigm. Participants read sentences word-by-word, with each word presented for 150 ms and no blank screen in-between words. After sentence reading, there was a short distraction task, where a list of four numbers was presented, followed by a single number written in letters. Participants had to decide whether the last number was present in the list. After their response, they repeated aloud the sentence they had just read as they remembered it (see Figure 1). This paradigm was shown to lead to sentence production from conceptual memory of the sentence just read, rather than from verbatim memory (Potter and Lombardi 1990; Lombardi and Potter 1992; Potter and Lombardi 1998; van de Velde and Meyer 2014). We thus considered it to be a suitable paradigm to elicit sentence production with constrained stimuli. Participants were asked to always utter full sentences, even if they did not remember one of the items they read, by using one of the other words they had been exposed to. These partly inaccurate sentences were also used as stimuli of interest. Audio was recorded in the scanner using an MRI-compatible microphone (Optoacoustics, FOMRI III) to code for the content of each sentence uttered.

**Figure 1:**
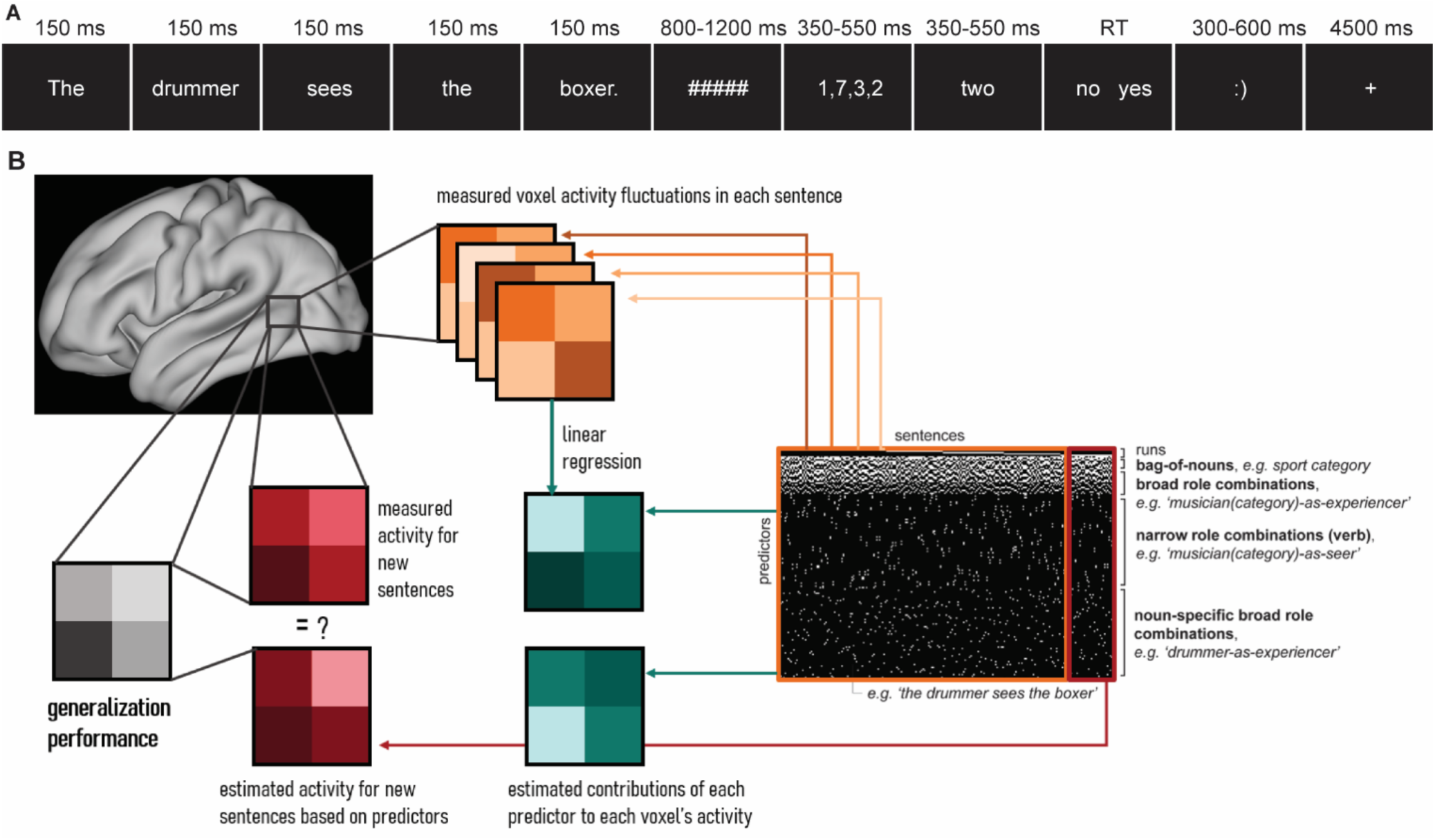
**A**: Experimental procedure including presentation times and jittering between events. The inter-trial interval ranged from .5 to 3.5 sec (mean = 0.64 sec). The sentence was first presented word-by-word. Then, as part of a distraction task, four number were presented, followed by a number written in letter. Participants had to indicate whether this latter number was present in the previous list of four numbers with yes or no. **B**: Visual representation of the encoding model procedure. The black and white matrix on the right contains all the predictors used in each encoding model of brain activity. Only a section of the predictors was used for each encoding model, as indicated by the examples. The white spots indicate that a sentence feature (e.g. semantic category for the bag-of-nouns) was present in a given sentence. The estimated contributions of each predictor (in green) to each voxel’s activity were calculated with linear regression of the voxel activity fluctuations in a training set of sentences (in orange). Activity for remaining test sentences (in red) was estimated based on the predictor estimates and then compared to the measured activity for new sentences. The comparison of estimated and measured activity led to a generalization performance measure (in grey) for each voxel.

The experiment was divided in 6 blocks of 60 sentences each. In 6 trials per block, a question was asked in place of producing a sentence. The questions asked about the identity of the agent or patient of the sentence just read (e.g. “Who kicked?” and a choice between “boxer” and “musician”). The questions were meant to have a measure of participants’ understanding of what they were reading. Twelve additional trials per run were filler sentences, leading to 42 sentences of interest per run. These sentences were selected to have a similar distribution of nouns, verbs and category combinations in each run.

### fMRI data acquisition

MR data were acquired in a 3T MAGNETOM PrismaFit MR scanner (Siemens AG, Healthcare Sector, Erlangen, Germany) using a 32-channel head coil. The MRI protocol included a T1-weighted MRI scan for anatomical reference and several fMRI scans. The T1-weighted scan was acquired in the sagittal orientation using a 3D MPRAGE sequence with the following parameters: repetition time (TR)/inversion time (TI) 2300/1100 ms, echo time (TE) 3 ms, 8° flip angle, field of view (FOV) 256 mm × 216 mm × 176 mm and a 1 mm isotropic resolution. Parallel imaging (iPAT = 2) was used to accelerate the acquisition resulting in an acquisition time of 5 min and 21 sec. Whole-brain functional images were acquired using a 3D EPI sequence following the implementation of Stirnberg et al. (Narsude et al. 2016; Stirnberg et al. 2017). This choice was motivated by the need to have a short TR to help separability of the tasks, while keeping a good sensitivity to internal brain structures and reducing the sensitivity to motion (multi-band sequences are more sensitive to motion and with poorer resolution in medial structures). These were the parameters: TR 700 ms, TE 33 ms, flip angle 15°, FOV 210 mm x 210 mm x 150 mm, voxel size 2.5 mm isotropic. Fieldmap images were also acquired to correct for distortions. We acquired 6 fMRI runs per participant.

### Behavioural analysis

The recording of each sentence was transcribed by a Dutch native speaker and rated for accuracy. We used two accuracy measures. One is standard accuracy for memory, where the sentence was rated to be correct if it was identical to the sentence that was just read. The other accuracy measure determined whether a sentence was suitable for the analysis, that is, whether the subject, object and verb were all part of the stimuli of interest, even if they did not match the sentence just presented. We decided to include these sentences in the analysis because we were primarily interested in sentence production, rather than correct memory. We also extracted the onset and offset times of the speech using Praat, after scanner noise removal. The onset and offset times were used for the timing of the production during fMRI analysis.

### fMRI preprocessing

Preprocessing was performed using *fMRIPrep* 20.2.6 (Esteban et al. 2018; Esteban et al. 2018 Dec 10), which is based on *Nipype* 1.7.0 (Gorgolewski et al. 2011; Gorgolewski et al. 2018). Details on the preprocessing are provided in the Supplementary Information.

The T1-weighted (T1w) image was corrected for intensity non-uniformity (INU). The T1w-reference was then skull-stripped and spatially normalized to MNI standard space (MNI152NLin2009cAsym). For each of the 6 BOLD runs per subject, the following preprocessing was performed. First, a reference volume and its skull-stripped version were generated using a custom methodology of *fMRIPrep*. A B0-nonuniformity map (or *fieldmap*) was estimated based on a phase-difference map. The *fieldmap* was then co-registered to the target reference run and converted to a displacements field map. Based on the estimated susceptibility distortion, a corrected reference was calculated for a more accurate co-registration with the anatomical reference. The BOLD reference was then co-registered to the T1w reference. Head-motion parameters with respect to the BOLD reference were estimated. The BOLD time-series were resampled into standard MNI space. Estimation of motion artifacts using independent component analysis (ICA-AROMA, Pruim et al. 2015) was performed on the *preprocessed BOLD on MNI space* time-series after removal of non-steady state volumes and spatial smoothing with an isotropic, Gaussian kernel of 6mm FWHM (full-width half-maximum). These noise-regressors were collected for nuisance regression in first-level analysis. Several confounding time-series were calculated based on the *preprocessed BOLD*: framewise displacement (FD), the derivative of the relative (frame-to-frame) bulk head motion variance (DVARS) and three region-wise global signals. Additionally, a set of physiological regressors were extracted to allow for component-based noise correction (*CompCor*, Behzadi et al. 2007). For anatomical CompCor, three probabilistic masks (CSF, WM and combined CSF+WM) are generated in anatomical space. The head-motion estimates calculated in the correction step were also placed within the corresponding confounds file.

### First-level analysis for sentence *t* map extraction

To extract beta and *t* maps on which encoding analyses were run, we ran a first-level analysis in SPM12 in Matlab 2021a. We ran a GLM on the preprocessed files in MNI space and we included confound regressors from fMRIPrep for DVARS, Framewise Displacement, 6 aCompCor parameters and 6 motion parameters. We also added the AROMA noise components computed in fMRIPrep as additional nuisance regressors, to perform non-aggressive denoising, that better accounts for motion-related noise. The design matrix included as condition regressors: one single regressor for all sentences read, one single regressor for the number task (from the presentation of the list of numbers to the response), one regressor for fillers, one regressor for question trials, and one regressors for all sentences that were erroneous. For the voxel-wise encoding model procedure, we included one regressor of interest per sentence, thus extracting about 40 beta maps per run (depending on the number of correct productions in each participant). For the voxel-wise encoding models during sentence comprehension, we used a very similar model with instead one regressor of interest modelling the time of sentence reading for each sentence separately, a single regressor for all produced sentences, as well as the number and filler regressors, extracting 48 beta maps per run, one for each sentence read (including question sentences). The average framewise displacement was 0.16 ± 0.006 across participants.

### Encoding model analyses

#### Encoding models

We compared encoding models that hinged on different aspects of the sentence (Fig. 1B). As introduced above, each sentence can be described in several ways: by its thematic roles, as well as by event information provided by the verb. We used contact verbs, that take Agent and Patient thematic roles, and perception verbs, that take Experiencer and Stimulus thematic roles (experiencer-subject psych verbs, Grimshaw 1990; Levin and Hovav 2005). We focused on encoding models that represent the sentence structure at different levels of (compositional) specificity (Table 2). Among these models of relational specificity, we used a bag-of-nouns model, which included the category of the nouns present in the sentence independently of their role in the sentence (4 of the 6 sentences in Table 2 would thus be deemed identical). This model had 2 parameters, one for each semantic category of the nouns. We also used a model encoding broad noun-role combinations across verbs (musician-as-agent), leading to 8 parameters (4 thematic roles x 2 noun categories). The agent/patient, stimulus/experiencer thematic roles were kept separate. In this way, we implicitly distinguished between event types (verb categories) as well, because in our stimuli agent/patient roles only occurred with contact verbs, and stimulus/experiencer roles with perception verbs. Finally, a narrow model targeted noun-verb specific combinations, leading to 64 parameters (2 semantic categories for nouns x 16 verbs x 2 thematic roles). These models thus characterized each sentence with different levels of specificity in the relations between nouns in the event.

**Table 2:**
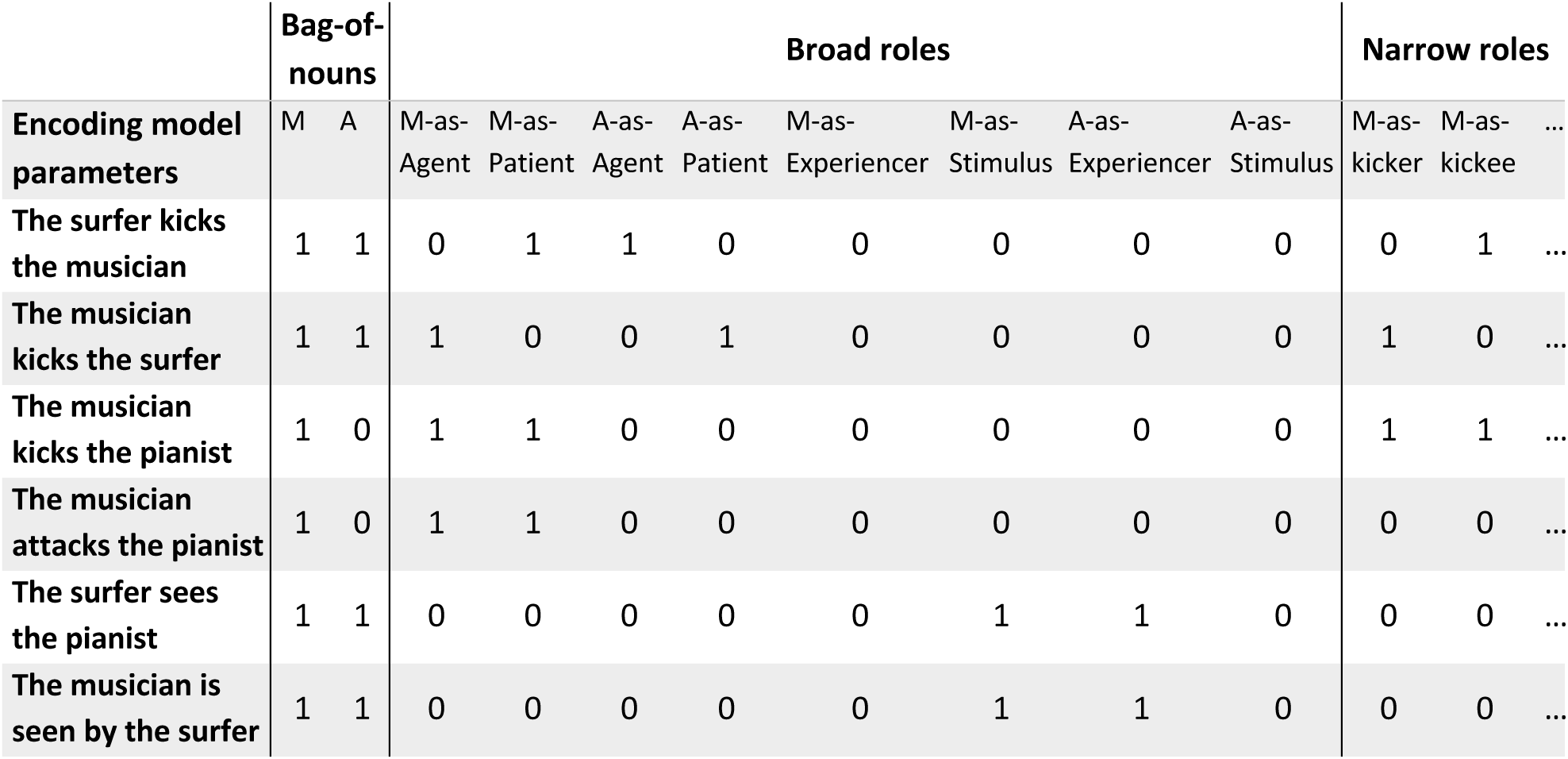
Encoding model parameters for the bag-of-nouns, broad roles and narrow roles models, with values for example sentences. M stands for semantic category “Musician”, A for semantic category “Athlete”. The narrow roles model had four columns per verb (two semantic categories, two thematic roles), leading to 64 parameters, of which only two are shown here.

#### Voxel-wise encoding model procedure

The voxel-wise encoding model analyses followed the procedure in Frankland and Greene (2020a). Some differences were unavoidable due to the different design and materials. The encoding models were run in Nilearn (Abraham et al. 2014; nilearn/nilearn 2022) on the beta maps for each sentence in a grey matter mask that included the cortex and subcortical structures relevant in production (caudate and putamen, hippocampus and cerebellum).

The encoding models were trained to predict the activity associated with each sentence in each voxel as a linear combination of sentence dimensions. The models were trained on 5 out of 6 runs and tested on the held-out sentences from the remaining run in a 6-fold cross-validation procedure. We extracted beta parameters per sentence descriptor per voxel per subject in a multiple regression of the sentence descriptors relevant for each encoding model. The model’s performance was then evaluated by comparing observed values with predicted values generated from the learned beta parameters for each voxel for the held-out sentences. The comparison was performed with z-scored squared differences between the predictions and observations in each run (leading to a predictions x observations squared matrix). We then averaged the on-diagonal elements (which correspond to correct mappings between predicted and observed sentences, i.e. predicted activity for sentence_i_ vs. observed activity for sentence_i_) and the off-diagonal elements (corresponding to incorrect mappings, i.e. predicted activity for sentence_i_ vs. observed activity for sentence_j_). Finally, we compared these averages: if the voxel encoded information on the sentence descriptors, the average for the correct mappings (difference between predicted and observed values) should be lower than the average for the incorrect mappings. The difference between correct and incorrect mappings was then averaged across the 6 runs for each voxel and multiplied by −1, so that informative voxels were represented as greater than zero. We then smoothed the maps at 8 mm FWHM, since the encoding model procedure was run on unsmoothed preprocessed data on each voxel separately.

#### Group-level whole-brain and small-volume correction analysis

To run group-level whole-brain analysis, we used a permutation procedure in pyMVPA (Stelzer et al. 2013). We created 100 permutations for each encoding model after permuting the sentence descriptors for each sentence. A cluster forming threshold was estimated via bootstrapping of the permutations (n=100 per model per participant), where permutation maps were randomly selected per subject 10,000 times, leading to 10,000 estimated group-average accuracy maps under the null hypothesis. For whole-brain analyses, we used a *p* < 0.005 voxelwise threshold, and a *p* < 0.05 cluster correction (False Discovery Rate, Benjamini-Hochberg procedure (Benjamini and Hochberg 1995)).

#### ROI analysis

For the exploratory analysis in several left-lateralized language regions, we extracted six masks for each region of interest using the Harvard Oxford atlas (Desikan et al. 2006), defining the LIFG *pars opercularis*, LIFG *pars triangularis* and left Precentral Gyrus using the corresponding labels available. The left anterior temporal lobe (LATL) mask included the left anterior superior and left anterior middle temporal gyri regions. The left posterior temporal lobe (LPTL) mask included the left posterior superior and middle temporal gyri as well as the left middle temporo-occipital cortex. The left temporo-parietal junction (LTPJ) mask included the left angular gyrus and left posterior supramarginal gyrus. We then isolated voxels in all ROIs that were found to have significant generalization performance for each model separately. We selected voxels independently for each participant, by selecting voxels with *t* > 2.4 (or *t* > 1.68 if no voxels survived the higher threshold) in all participants except for one held-out participant iteratively. We could then localize regions sensitive to each model independently for each participant and averaged each model’s performance in these regions. We then ran linear mixed-effects models with *lme4*, predicting models’ generalization performance with factors ROI, modality (production and comprehension) and model (bag-of-nouns, broad roles and narrow roles), and by-participant random slopes for ROI and class. We ran pairwise comparisons using *emmeans* (Lenth et al. 2022).

## Results

### Behavioural results

First, we determined whether participants suitably performed the distraction task, thus ensuring they were not actively rehearsing the sentence just read to prepare for production. All participants had above chance accuracy in the distraction task (mean = 0.91, SE = 0.01, with average reaction time 451.1 ms, SE = 16.7, Fig. 2). Next, we assessed the performance on the main sentence recall task. Performance varied consistently across participants (Fig. 2). Some participants exhibited relatively poor memory for the sentence just read, while others had very high accuracy (mean = 0.77, SE = 0.02). However, we were not just interested in the production of sentences identical to the ones just read, but we also considered as acceptable sentences that were constituted by words that were part of the stimulus set (Table 1). These sentences increased the separability between stimuli read during comprehension and spoken during production (23% of acceptable sentences were different from previously read sentences). The number of ‘acceptable’ sentences was higher than ‘accurate’ sentences across participants (mean = 0.93, SE = 0.007), which led to a high number of stimuli and served as confirmation that participants could perform the task appropriately. Incorrect sentences included words that were not part of the stimuli (e.g. uttering “man” instead of “musician”), sentences with different structure, or sentences that did not include two participants and a verb.

**Figure 2:**
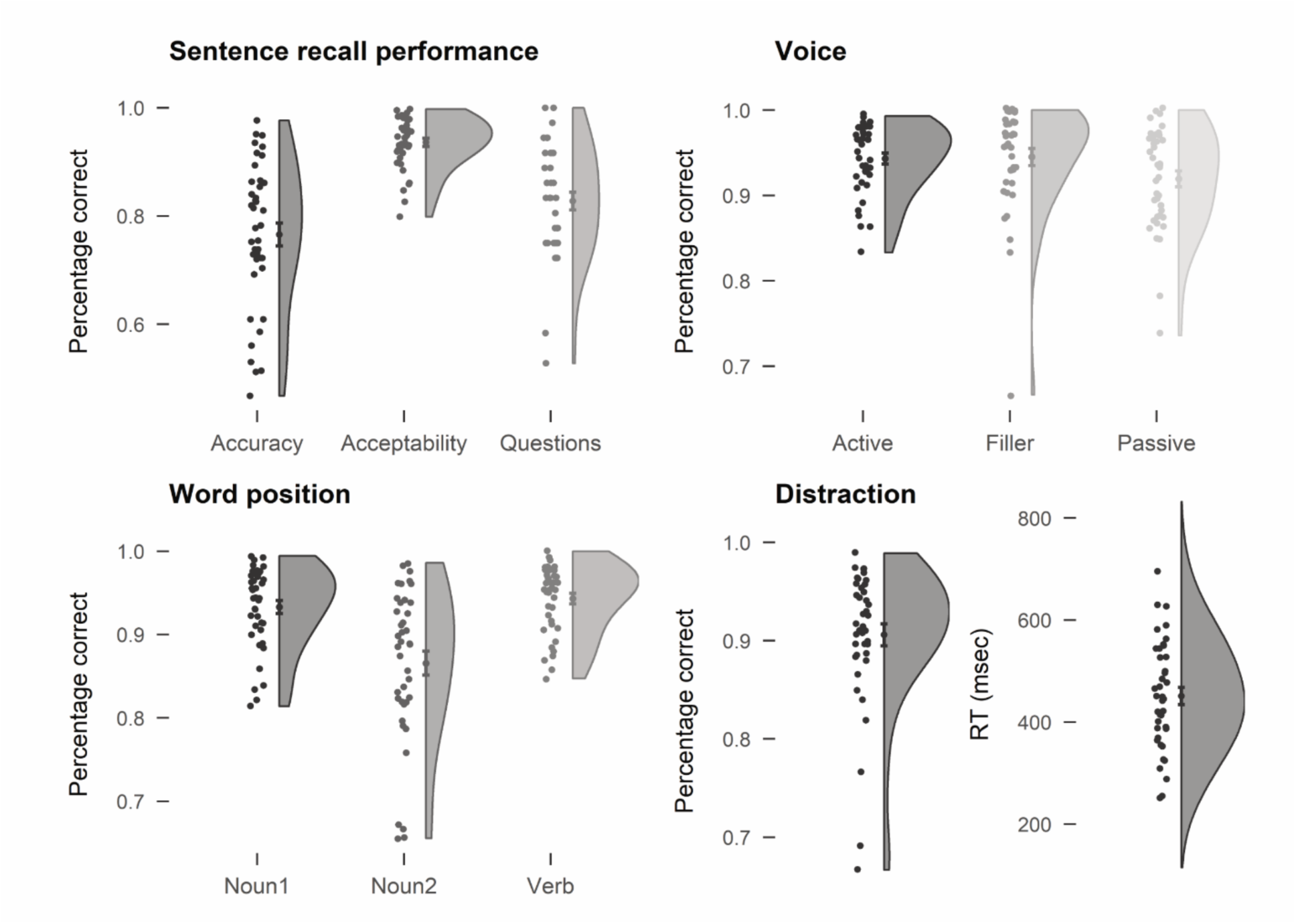
Summary of the behavioural performance. Points on the left of the violin plot represent individual participants’ performance. Points on the violin plot represent the mean, error bars standard error of the mean. Voice performance shows how many acceptable sentences were produced in each condition. Word position reports accuracy values in the memory of words for each position in a sentence.

We also determined what parts of the sentence were more prone to errors and, therefore, possibly encoded less strongly. Verbs had the highest accuracy (mean = 0.94, SE = 0.006), followed by the first noun (mean = 0.93, SE = 0.007; verb vs. first noun: *t* = 2.35, *p* < 0.05), while the second noun was the most likely to be forgotten (mean = 0.86, SE = 0.014; *t*s > 7.7, *p* < 0.0001). Errors were more likely to happen in passive relative to active sentences (accuracy, active = 0.83, SE = 0.017; passive = 0.72, SE = 0.025; *t* = 9.8, *p* < 0.0001; acceptability, active = 0.94, SE = 0.006; passive = 0.92, SE = 0.009; *t* = 3.8, *p* = 0.0005). Finally, we checked the performance during question trials, to make sure participants were correctly parsing the sentences. Performance was overall good (mean = 0.83, SE = 0.016), but one participant with accuracy around chance level was excluded (accuracy = 0.53).

### Specificity of event encoding: whole-brain results

#### Sentence production

First, we searched the whole brain for clusters encoding thematic roles at different levels of specificity. We compared a bag-of-nouns model to a broad roles model and a narrow roles model. The bag-of-nouns model was defined based on the presence of music and sport actors (by category) in the sentence, taking the category of all words in italics independently of position and thematic role: “the *violinist* attacks the *surfer*”, categorized as *musician* and *athlete* present. The broad roles model was defined based on the presence of a semantic category and its thematic role: musician-as-agent/stimulus/patient/experiencer: “the *violinist* attacks the *surfer*” was encoded as *musician-as-agent*, *athlete-as-patient*. The narrow roles model was defined based on the presence of a semantic category together with the thematic role specific to the verb: “the *violinist* attacks the *surfer*” encoded as *musician-as-attacker* and *athlete-as-attackee*, using the specific verbs instead of their semantic categories. Since thematic roles agent/patient and stimulus/experiencer are defined based on the type of verb, the broad roles model encoded thematic roles across verbs, but not independently of verb categories (i.e. contact or perception verbs). The narrow roles model instead included idiosyncrasies in verb meaning. In all these three models the event participants were not defined individually, but by their semantic categories.

All encoding models were significant in many fronto-temporal clusters (Fig. 3, see Supplementary Table 1 for peaks). The bag-of-nouns model led to the most widespread clusters, in large parts of left posterior superior, middle and inferior temporal gyri, extending into inferior parietal lobe, including supramarginal and angular gyri, and to inferior and middle frontal gyri (LIFGtri and LIFGoper), as well as precentral gyrus and supplementary motor cortex. There were additional clusters in the left and right middle occipital cortex, anterior cingulate cortex and medial prefrontal cortex, and right STG, ITG and IFG. The broad roles model had largely overlapping clusters in left and right STG, left and right supramarginal and angular gyri, LIFGtri and LIFGoper, bilateral precentral gyri middle frontal gyrus and supplementary motor cortex, with an additional cluster in the right cerebellum (lobule VII). The narrow roles model for verbs instead was more limited, with a cluster in left posterior STG extending into supramarginal and angular gyri, and a cluster in precental gyrus and LIFGoper extending to LIFGtri and *pars orbitalis*. There were additional clusters in the supplementary motor cortex, mid cingulate cortex, right STG and posterior MTG, right insula and right putamen. All models thus seemed to be centred in language-relevant areas, and to largely overlap in their distribution.

**Figure 3:**
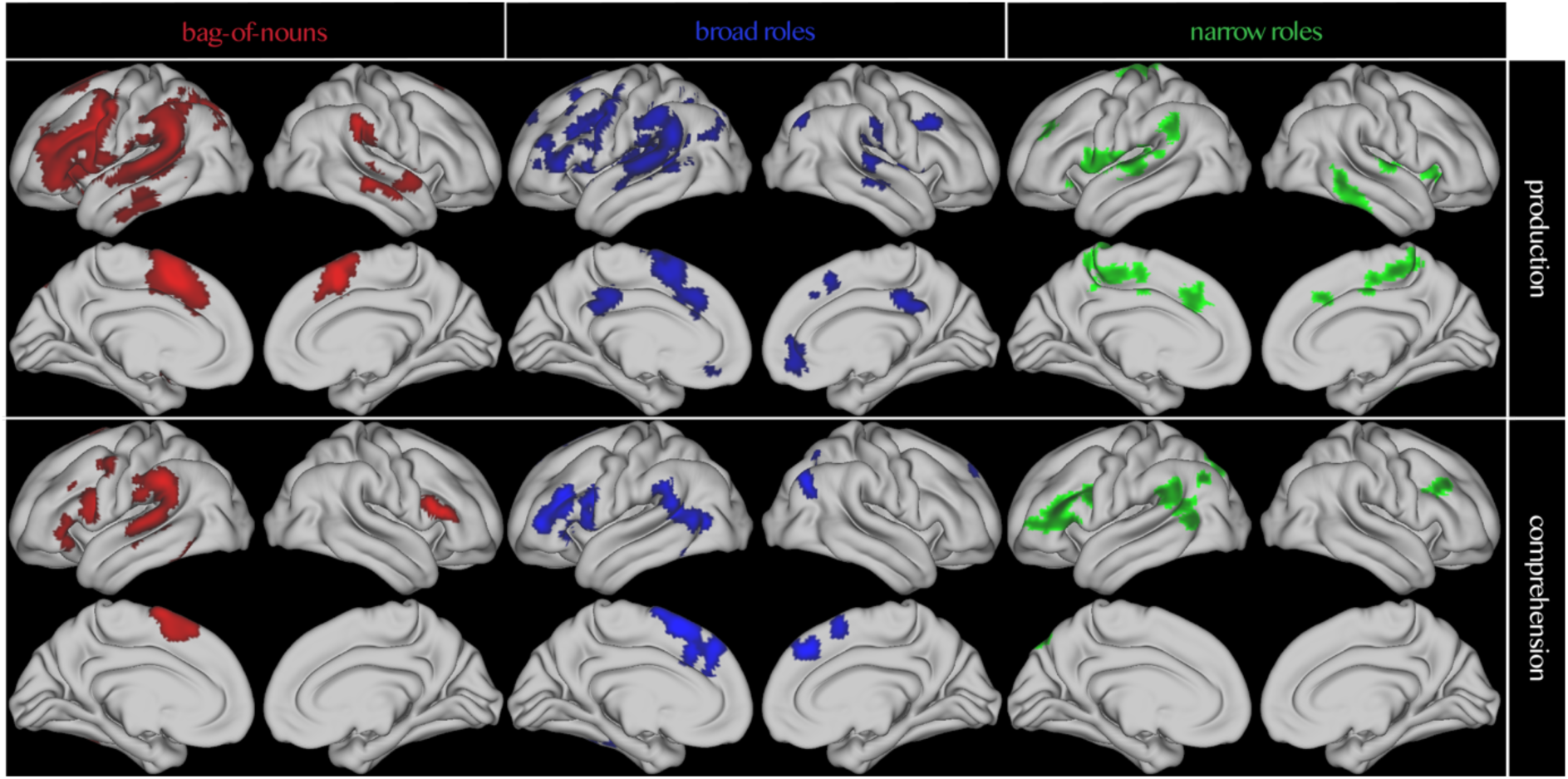
Whole-brain results for the generalization performance of the bag-of-nouns model, broad roles and narrow roles model in production and comprehension. All results are thresholded at p < 0.005, p < 0.05 FDR cluster corrected.

#### Sentence comprehension

Since a previous study with a similar approach but different design and stimuli had found a compositional network in sentence comprehension to be centred in the lmSTC, amPFC and hippocampus (Frankland and Greene 2020a), we asked whether the difference in results could be due to language modality (note that an ROI analysis on those three regions shows different results from Frankland and Greene (2020), presented in the Supplementary Information). We therefore ran the same whole-brain encoding analysis during the rapid comprehension of the sentences, preceding the distraction task. All models yielded significant clusters in the temporal parietal junction and posterior inferior frontal gyrus, with few differences among models (see Fig. 3 and Supplementary Table 2 for specific clusters and peaks). It should be noted that this study was designed primarily for sentence production, and that participants had a very short time to process these sentences (0.75-1.05 s, followed by 0.8-1.2 s fixation), so it is not surprising to find smaller effects in comprehension. Overall, the comprehension whole-brain results appeared to be similarly distributed as the production results.

### Are any clusters specific to a level of relational specificity?

The whole-brain results, therefore, suggest that both specific and abstract roles are encoded during sentence production across the language network. We then asked whether any regions showed increased generalization performance for the broad and narrow roles over the bag-of-nouns model. We ran a whole-brain paired t-test in SPM for the generalization performance of each model. We found no clusters showing significantly better generalization for any model. This lack of specificity for a single model over other models, or improved performance for one of the relational models over the bag-of-nouns model, could suggest that the relational models are not actually encoding composed semantic structures, but may be just capitalizing on the information on semantic categories that was still present across several more specific predictors, similarly to the bag-of-nouns model. Therefore, we tested whether the relational models had better performance than the bag-of-nouns model in an ROI analysis of regions of the language network. To do this, we parcellated this left-lateralized network in six regions: LATL, LPTL, LTPJ, LIFGoper, LIFGtri, LPrecentral Gyrus. We selected voxels in each participant based on model performance in all other participants. We then ran a mixed-effects model with a three-way interaction between model, modality and ROI (Fig. 4). We found a main effect of model (χ^2^ = 17.4, *p* < 0.002), ROI (χ^2^ = 30.5, *p* < 0.0001) and modality (χ^2^ = 3.9, *p* = 0.046), but no interaction. Performance was slightly higher in production (*t* = 1.97, *p* = 0.057), which is probably due to production having a stronger focus in this design. The narrow roles model performed significantly better than the bag-of-nouns and broad roles models (*t*s > 2.7, *p* = 0.023), confirming that modelling relational structure improves performance and that the generalization performance in the more complex models was not just capitalizing on the categorical information present in the anrrow roles model. Performance was significantly lower in the LATL than all other areas except for the LPTL (ts > 3.1, ps < 0.035), but not different among other models. Pairwise comparisons indicated that the narrow roles model performed better than the bag-of-nouns model in the LIFGtri and LTPJ in production, and in the LIFGtri, LIFGoper and LTPJ in comprehension. Note that the better performance of the narrow roles model together with the smaller clusters in the whole-brain analysis suggests that the narrow roles model had larger variance in its accuracy (due to the larger number of parameters relative to the number of training sentences), leading to relatively less power and smaller clusters in the whole-brain analysis.

**Figure 4:**
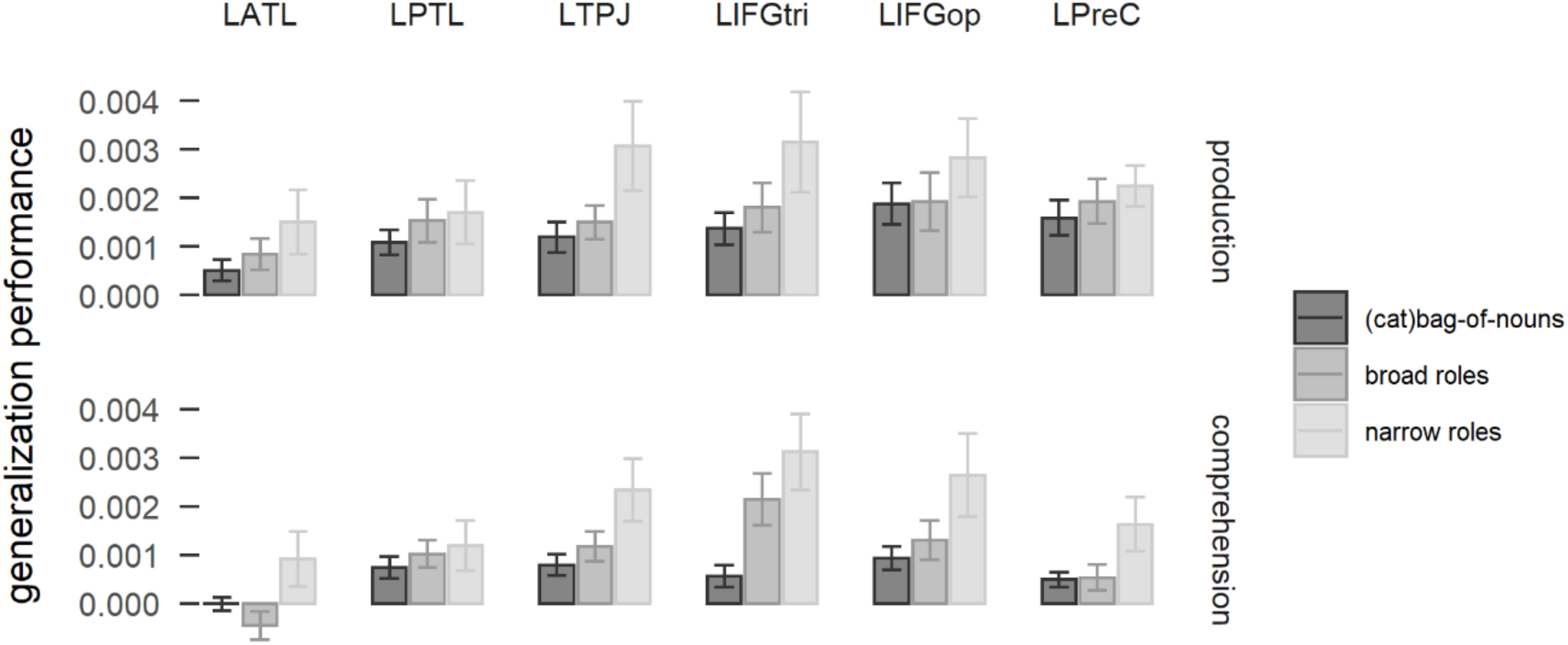
Generalization performance for the bag-of-nouns model (that separates the semantic categories of musician and athlete), the broad roles model and the narrow roles model, in six brain regions. LATL: left anterior temporal lobe. LPTL = left posterior temporal lobe. LTPJ: left temporo-parietal junction. LIFGtri: pars triangularis of the left inferior frontal gyrus. LIFGop: pars opercularis of the left inferior frontal gyrus. LPreC: left precentral gyrus.

### Is narrow role model performance driven by the more specific verb encoding?

The narrow roles model performed better than the bag-of-nouns model and the broad roles model in the network obtained by the whole-brain analysis. This raises the question if the improved performance is simply due to the more specific modelling of verb meaning in the narrow roles model, relative to the absence of verb meaning in the bag-of-nouns model and the types of thematic roles, which corresponded to the verb semantic categories, in the broad roles model. We therefore tested whether the narrow roles model showed improved performance relative to a non-relational bag-of-verbs model (with separate predictors for each verb, but not in combination with thematic roles), in a linear mixed-effects model with model, ROI and modality as fixed effects (Fig. 5). We found an interaction between model and ROI (χ^2^ = 12.1, *p* < 0.0005) and a trend for an interaction between modality and ROI (χ^2^ = 9.2, *p* = 0.09). Pairwise comparisons indicated that the relational information improved performance only in the LIFG in production (both LIFGtri and LIFGoper, *p* < 0.05), while there were no significant differences between models in the other ROIs and in comprehension. The results suggest that the addition of specific verb predictors (as opposed to categorical ones) improved generalization performance without relational information, but relational information was still driving improved generalization in the LIFG in production.

**Figure 5:**
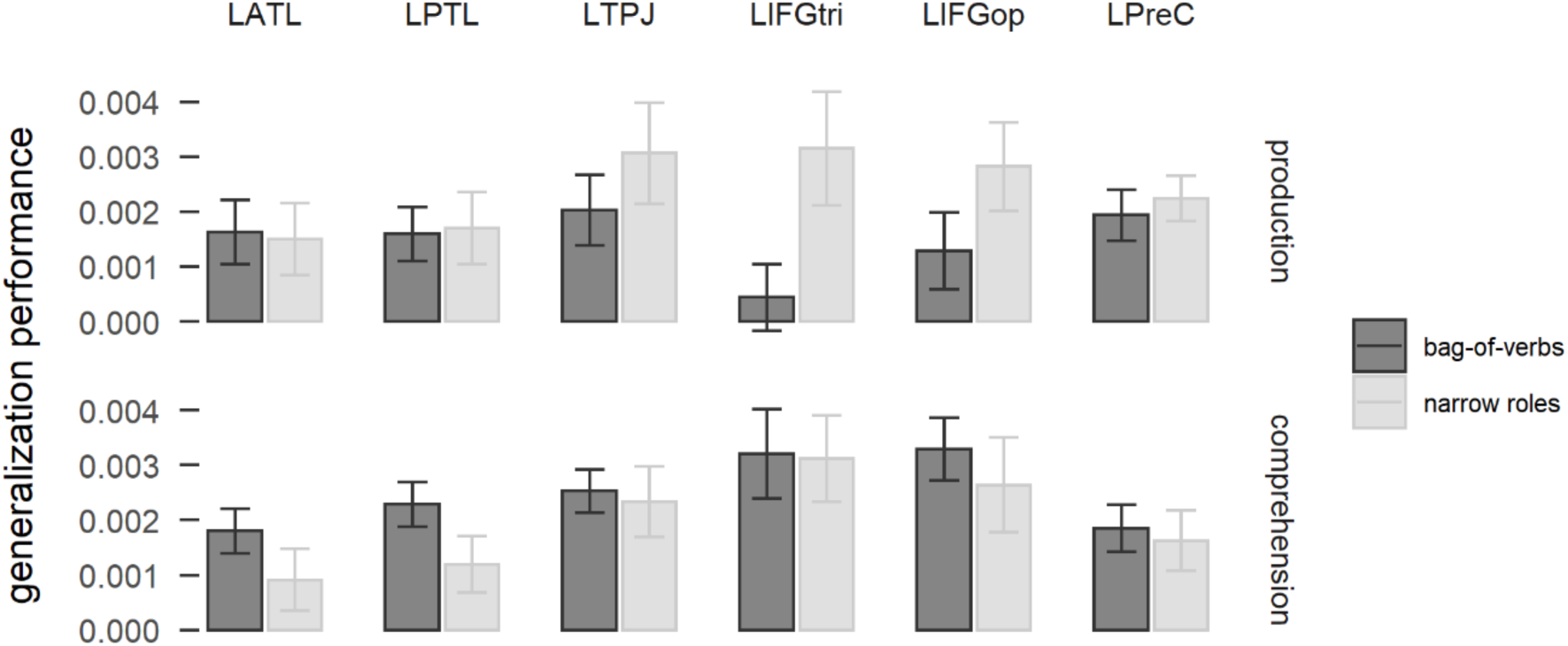
Generalization performance for the bag-of-verbs model that includes of all verbs and the narrow roles model, in six brain regions. LATL: left anterior temporal lobe. LPTL = left posterior temporal lobe. LTPJ: left temporo-parietal junction. LIFGtri: pars triangularis of the left inferior frontal gyrus. LIFGop: pars opercularis of the left inferior frontal gyrus. LPreC: left precentral gyrus.

### Does noun-specific encoding similarly improve generalization performance?

The high performance for verb-specific information raises the question if specific-noun information may similarly provide a better fit for activation patterns than the categorical bag-of-nouns model used so far. All the analyses described so far characterized nouns in terms of their thematic category (music vs. sport). Our decision to use categories was motivated by the preference for more varied production output and less repetition. Therefore, we ran a follow-up analysis to determine whether the specific noun meaning affected the way the event was encoded in the ROIs. We tested if a bag-of-nouns model including all the nouns, not merged in semantic categories, improved performance over the categorical bag-of-nouns. Additionally, we tested an encoding model where each noun was associated to a broad thematic role related to the verb category, i.e. noun-specific broad roles model (contact or perception) (e.g. “the violinist attacks the surfer” is encoded as *violinist-as-agent*, *surfer-as-patient*). We then determined the generalization performance of these models in the network parcels (Fig. 6).

**Figure 6:**
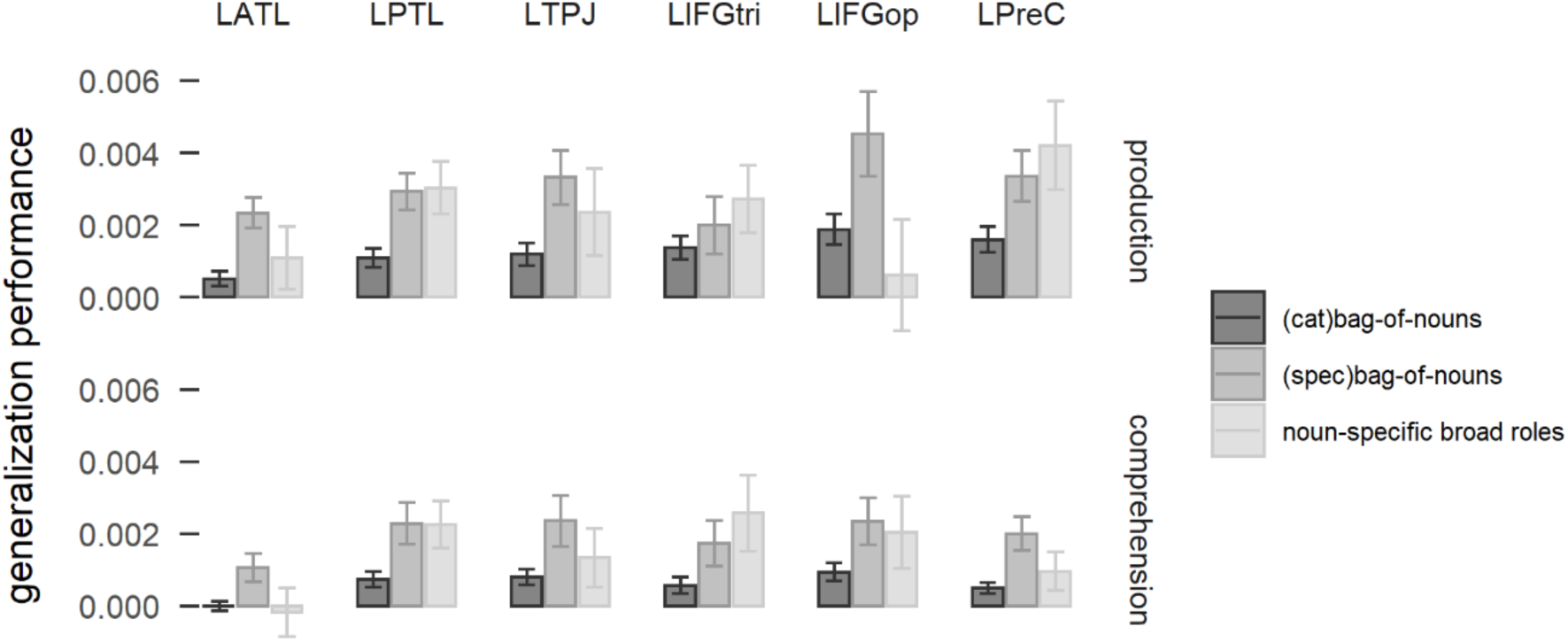
Generalization performance for the categorical bag-of-nouns model (that separates the semantic categories of musician and athlete), the specific bag-of-nouns model and the noun-specifical broad roles model, in six brain regions. LATL: left anterior temporal lobe. LPTL = left posterior temporal lobe. LTPJ: left temporo-parietal junction. LIFGtri: pars triangularis of the left inferior frontal gyrus. LIFGop: pars opercularis of the left inferior frontal gyrus. LPreC: left precentral gyrus.

We found a significant effect of ROI (χ^2^ = 19.4, *p* < 0.002), model (χ^2^ = 21.4, *p* < 0.0001), an interaction between ROI and model (χ^2^ = 23.5, *p* < 0.009) and an interaction between model, ROI and modality (χ^2^ = 18.5, *p* = 0.046). The specific bag-of-nouns model had overall better performance than the categorical bag-of-nouns model (estimate = 0.0016, *t =* 4.6, *p* = 0.001). The noun-specific broad roles model also had marginally better performance than the categorical bag-of-nouns model (estimate = 0.001, *p* = 0.098). There were no significant differences between the noun-specific broad roles model and the specific bag-of-nouns model in different ROIs, with the exception of the LIFGoper that showed significantly better performance for the specific bag-of-nouns model in production (estimate 0.004, *p* < 0.0001). The inclusion of individual noun predictors therefore improves encoding performance, relative to broad semantic categories. The addition of broad thematic roles does not significantly improve generalization performance. Relational information, therefore, seems to improve generalization performance relative to verb-specific encoding but not noun-specific encoding. The current design does not allow to test a relational model that contains both noun and verb-specific thematic information, since not all possible noun-verb combinations were produced by participants (16 nouns x 16 verbs x 2 thematic roles = 512 predictors for an average of 240 ± 10 sentences per participant).

## Discussion

In an fMRI study of compositional processing in sentence production, we used voxel-wise encoding models to investigate how the brain builds compositional meaning during sentence production. We looked for regions encoding sentence meaning with three models. A bag-of-nouns model encoded the semantic categories of nouns present in a sentence, either musicians or athletes. A broad roles model encoded the semantic category of the noun in combination with its thematic role, agent and patient or experiencer and stimulus, based on the presence of verbs defining a contact event or a perception event. A narrow roles model encoded verb-specific relational structure in combination with the noun semantic categories. Whole-brain analyses revealed that a left fronto-parieto-temporal network encoded semantic structures at different levels of specificity, both during the comprehension and the production of sentences. This network overall encoded specific relational information, as modelled by the narrow roles model, more strongly than the other models. There was also a preference for noun- and verb-specific information over semantic categories. Overall, our results suggest a distributed encoding of sentence meaning in the language network.

The whole-brain results showed that the relational structure of the sentence was encoded in several areas of the perisylvian language network. The different models had some differences in their responses, but there was considerable overlap. This network included mid-to-posterior superior temporal cortex, temporo-parietal junction, and inferior and middle frontal gyri extending into the precentral gyrus, with supplementary motor area. This network is in line with the brain regions found to represent distinctions for thematic categories (i.e. music vs. sport categories in this context) in a previous study, which peaked in the temporoparietal junction and included precentral gyrus and parietal areas (Xu et al. 2018). Thematic errors in aphasia are also associated with lesions in the temporo-parietal junction and white matter tracts between the temporal pole and precentral cortex (Schwartz et al. 2011; Schwen Blackett et al. 2022). A previous MEG study also found that compositional sentence meaning is encoded in interaction between temporal and inferior frontal cortex in a network that reminds of the current one (Lyu et al. 2019). In the current study, models with increasing specificity (i.e. broad roles and even more narrow roles) were characterized by less extensive clusters. One reason for this difference with the more extensive bag-of-nouns network is that the narrow roles model had many more parameters to model, and, as a consequence, fewer sentences on which to train. Therefore, the more specific models may have had less reliable estimates, leading to reduced power and fewer and smaller clusters (although note that the generalization performance for more complex models was overall higher).

An ROI analysis in a few regions of the language network (e.g. Brennan et al. 2016; Hu et al. 2022) showed that the narrow roles model had higher generalization performance than the broad roles and the bag-of-nouns model overall in this network, confirming that the performance of the narrow roles model was not simply due to the presence of semantic categories in the model. However, the improved performance in many regions seemed to be driven by the encoding of verb-specific information rather than specifically noun-verb combination indicating potential compositional sentence meaning. The only exception were the LIFGtri and LIFGoper parcels, which showed higher performance for the narrow roles model than the individual verb model (and less strongly the LTPJ) in production. Therefore, it seems that all of the regions in the network encoded the meaning of the different parts of the sentence, and in particular noun semantic categories, verb identity and thematic roles, with the LIFG additionally shown to encode event-specific relational structure. The LIFG is a key language region that has been often implicated in syntactic processing and semantic unification (Bornkessel-Schlesewsky and Schlesewsky 2013; Hagoort 2013; Lyu et al. 2019; Hagoort 2020). Here, it is interesting to note that the LIFGoper was found to encode non-relational noun-specific information but also relational verb-specific information, while the LIFGtri encoded relational and non-relational specific information (while being less sensitive to semantic categories). This tentatively suggests meaningful interactions between these regions in keeping relevant information in working memory for composition, as well as for phonological encoding (Xiang et al. 2010; Conner et al. 2019; Morgan et al. 2024).

An additional follow-up analysis showed that, while semantic categories were successfully encoded, there was a preference for noun specific identity, over semantic categories. On the one hand, the significant performance for semantic categories shows that the brain encodes meaning along semantic features, as previously shown (Xu et al. 2018; Elli et al. 2019). On the other hand, it is clear that the specific features of each noun and verb are decodable more successfully from brain activity, even if it means that there are fewer observations per item. This finding additionally raises the question of how an encoding model with a combination of both noun- and verb-specific identity would have done. It was not possible to test this model here due presence of too many parameters for a fully specified relational model relative to the number of sentences produced.

A previous study focused on compositional sentence meaning in comprehension using fewer nouns and verbs and modelling relational structure using noun- and verb-specific identities (Frankland and Greene 2020a). They found evidence for the lmSTC encoding abstract relational structure (broad roles), the amPFC encoding event specific relational structure (narrow roles) and the hippocampus encoding event specific relational structure in a pattern separation fashion (Frankland and Greene 2020a). Our whole-brain results were not significant in these regions, suggesting either that the inclusion of noun-specific identity may have led to the finding of additional regions, or that the network encoding relational structure may be more distributed. Specific analyses into these regions (Supplementary Information) showed that only the lmSTC encoded relational structure among these ROIs in our study. The lower performance in the hippocampus and amPFC in the current study may be due to the poorer signal-to-noise ratio in some regions, as a consequence of the very fast TR that reduced signal in subcortical regions and perhaps in regions prone to signal dropout. Frankland and Greene (2020a) additionally noted that relational information in combination with semantic categories was not encoded in the three ROIs, but was found instead in other brain regions in the lateral prefrontal cortex and LSTG/IFG, in agreement with our current findings of a larger network involved in semantic composition.

The results of the previous study suggest that our stimulus set likely limited the encoding potential we could explore. In fact, we focused on semantic categories, and only included verb-specific role combinations (‘musician(category)-as-attacker’), but not noun-verb specific combinations (e.g. ‘violinist-as-attacker’), since the stimuli each participant produced did not span all of the combinations. It is therefore possible that specific noun-verb combinations may have improved generalization performance, under the assumption that the verb-specific combination could only benefit the encoding model if bound to specific nouns rather than semantic categories. This explanation is also supported by the high generalization performance for specific noun and verb models in the parcels. Generalization performance was higher when the specific noun information was included in the model, relative to the semantic categories, suggesting that studies investigating noun-role combinations benefit from the inclusion of specific information (although modelling specific noun and verb combinations requires a limited range of nouns and verbs, largely reducing the variability in the stimulus set which possibly has consequences for ecological validity). Future studies may be able to further uncover the neural encoding of compositional sentence meaning in production and comprehension.

It is important to note that differences in results between our study and the previous comprehension study is unlikely to be due to modality (i.e. production vs. comprehension), since the whole-brain results in comprehension in this study closely resembled the production results. The ROI results in comprehension were instead less definitive, with similar regions to production encoding narrow roles more strongly, but no regions showing a preference for narrow roles over simple verb identity. Since this study was designed specifically for production and the presentation of sentences was very short in comprehension, we refrain from interpreting non-significant results in comprehension, as it is impossible to know if they are due to lack of power or effective differences in encoding performance.

After taking into consideration evidence from follow-up analyses, it seems that the LIFG alone convincingly encoded relational structure in sentence production. Therefore, this study provides evidence for the encoding of compositional representations during sentence production, despite incremental message generation. We found neural representations for compositional structure in a sentence recall task that highly differs from the scene description experiments with which message generation in production is usually studied (Konopka and Brown-Schmidt 2014). The task used here is thought to resemble sentence production starting from a conceptual representation built during an initial rapid comprehension of the sentence^1^ (Potter and Lombardi 1990; Lombardi and Potter 1992; Potter and Lombardi 1998), and is thus more similar to the conditions with which compositional processing is usually studied in sentence comprehension (e.g. Frankland and Greene 2015; Frankland and Greene 2020a; Arana 2022), relative to scene description experiments. Therefore, even though the conceptual representation of a sentence can be formed incrementally, with an event type (i.e. verb) possibly selected after speech onset (e.g. Momma et al., 2016), the words of the sentence are successfully composed into a specific relational structure that is decodable from brain activity. Although our results are not informative about timing, they show that rich semantic structure can be decoded from neural activity in production despite its incrementality, possibly because the rapid and spontaneous binding of thematic roles to event participants applies not only to scene processing but also to conceptualization in message generation (Hafri et al. 2013; Hafri et al. 2018).

In addition, the evidence for the brain encoding event relations in both abstract and specific terms, especially as highlighted by the whole-brain results, but also as shown by the brain parcels, supports previous findings of compositional semantic representations being encoded in different forms (i.e. abstract and verb-specific) across brain regions, suggesting that the brain represents *who did what to whom* with flexible representations (Frankland and Greene 2015; Frankland and Greene 2020a). The brain thus seems to compose meaning based on several aspects of the sentence that are encoded simultaneously in a large network. The identity of the participants of the event are encoded generally along semantic features as well as separately, but also bound to their thematic role and the specific event they are associated to. The current results additionally suggest the same set of regions can carry information about several levels of specificity.

Finally, we found widespread whole-brain results for several levels of specificity in relational structure, but the follow-up analyses suggest that the LIFG was most convincingly encoding relational information above word information. Therefore, while we find sensitivity to sentence meaning more generally throughout the language network, it seems there is a specialization for the more specific compositional processes, although with differences in the specific regions across studies (cf. Frankland and Greene 2020a). Other studies found similar distributed representations for several aspects of sentence comprehension as well as production. For example, many regions in the language network are sensitive to both syntactic and lexical structure (Fedorenko et al. 2012; Blank et al. 2016; Fedorenko et al. 2020; Shain et al. 2021; Hu et al. 2022). A very large network responded to constituent size in both sentence production and comprehension (Giglio et al. 2022). Similarly, the meaning of words in multiple positions in a sentence (e.g. subject, object) was found to be encoded in frontal, inferior parietal and temporal regions (Anderson et al. 2019). This pattern of results is found in univariate as well as multivariate studies. The same linguistic representations thus seem to be encoded in many brain regions, and multiple levels of linguistic representations are seen to be encoded within the same brain region. The growing evidence for distributed responses stresses the importance of functional connectivity between brain regions in the interpretation of fMRI results (Friston 2002). In fMRI, we are characterizing brain activity with a very low temporal resolution (e.g. here we take the average brain activity over the course of sentence production, i.e. a few seconds, relative to more dynamic processes, Morgan et al. 2024). In the faster pace of neural activity, many brain regions may have been active in a rapidly firing network that builds these representations in integration across several regions, even if we are only able to capture the static distributed pattern (Thiebaut de Schotten and Forkel 2022). This is a view that is becoming more emergent in recent cognitive neuroscience (see also Hagoort 2019), and is especially corroborated by the many MEG studies that show fast connectivity between brain regions during sentence processing, in a dynamic network that cannot be easily characterized with fMRI (e.g. Schoffelen et al. 2017; Hultén et al. 2019; Lyu et al. 2019; Huizeling et al. 2022). An appropriate characterization of compositional sentence processing will therefore benefit from investigations with more time-resolved methods.

### Conclusions

Overall, we found that an extensive fronto-parieto-temporal network encoded sentence meaning during sentence production. A similar network was identified during the comprehension of the same sentences, pointing to a shared network for sentence meaning in sentence production and comprehension. While this network encoded event-specific information more strongly than the identity of the actors in the event, only the LIFG was seen to encode relational information over and above verb identity.

## Acknowledgements

We thank Steven Frankland and two anonymous reviewers for comments on an earlier version of the manuscript. We also thank Birgit Knudsen, Iris Schmits and Marjolijn Dijkhuis for support with coding of the production recordings. This work was supported by the Max Planck Society. P.H. was supported by the NWO Grant Language in Interaction, grant number 024.001.006.

## Data Availability

The data will be made available on the Radboud Data Repository (https://data.ru.nl/) upon publication.

## Footnotes

1 Note that it is unlikely that the representation we decoded was in effect from the comprehension of the sentence, since production followed a distraction task, and the produced sentences differed from the sentences comprehended on 23% of trials.

