## Supplementary Information for "Neural Encoding of Semantic Structures During Sentence Production"

### fMRIprep preprocessing pipeline

#### Anatomical data preprocessing

The T1-weighted (T1w) image was corrected for intensity non-uniformity (INU) with N4BiasFieldCorrection, distributed with ANTs 2.3.3 (Avants et al. 2008) and used as T1w-reference throughout the workflow. The T1w-reference was then skull-stripped with a *Nipype* implementation of the antsBrainExtraction.sh workflow (from ANTs), using OASIS30ANTs as target template. Brain tissue segmentation of cerebrospinal fluid (CSF), white-matter (WM) and gray-matter (GM) was performed on the brain-extracted T1w using fast (FSL 5.0.9, Zhang et al., 2001). Brain surfaces were reconstructed using recon-all (FreeSurfer 6.0.1, Dale et al., 1999), and the brain mask estimated previously was refined with a custom variation of the method to reconcile ANTs-derived and FreeSurfer-derived segmentations of the cortical gray-matter of Mindboggle (Klein et al. 2017). Volume-based spatial normalization to MNI standard space (MNI152NLin2009cAsym) was performed through nonlinear registration with antsRegistration (ANTs 2.3.3), using brain-extracted versions of both T1w reference and the T1w template.

#### Functional data preprocessing

For each of the 6 BOLD runs per subject, the following preprocessing was performed. First, a reference volume and its skull-stripped version were generated using a custom methodology of *fMRIPrep*. A B0-nonuniformity map (or *fieldmap*) was estimated based on a phase-difference map. The *fieldmap* was then co-registered to the target EPI (echo-planar imaging) reference run and converted to a displacements field map. Based on the estimated susceptibility distortion, a corrected EPI (echo-planar imaging) reference was calculated for a more accurate co-registration with the anatomical reference. The BOLD reference was then co-registered to the T1w reference using bbregister (FreeSurfer) which implements boundary-based registration (Greve and Fischl 2009). Co-registration was configured with six degrees of freedom. Head-motion parameters with respect to the BOLD reference (transformation matrices, and six corresponding rotation and translation parameters) are estimated before any spatiotemporal filtering using mcflirt (FSL 5.0.9, Jenkinson et al., 2002). The BOLD time-series were resampled into standard space, generating a *preprocessed BOLD run in MNI152NLin2009cAsym space*. First, a reference volume and its skull-stripped version were generated using a custom methodology of *fMRIPrep*. Automatic removal of motion artifacts using independent component analysis (ICA-AROMA, Pruim et al. 2015) was performed on the *preprocessed BOLD on MNI space* time-series after removal of non-steady state volumes and spatial smoothing with an isotropic, Gaussian kernel of 6mm FWHM (full-width half-maximum). The "aggressive" noise-regressors were collected for nuisance regression in first-level analysis. Several confounding time-series were calculated based on the *preprocessed BOLD*: framewise displacement (FD), the derivative of the relative (frame-to-frame) bulk head motion variance (DVARS) and three region-wise global signals. FD was computed using two formulations following Power (absolute sum of relative motions, Power et al. (2014)) and Jenkinson (relative root mean square displacement between affines, Jenkinson et al. (2002)). FD and DVARS are calculated for each functional run, both using their implementations in *Nipype* (following the definitions by Power et al. 2014). Additionally, a set of physiological regressors were extracted to allow for component-based noise correction (*CompCor*, Behzadi et al. 2007). Principal components are estimated after high-pass filtering the *preprocessed BOLD* time-series (using a discrete cosine filter with 128s cut-off). For anatomical CompCor, three probabilistic masks (CSF, WM and combined CSF+WM) are generated in anatomical space. The head-motion estimates calculated in the correction step were also placed within the corresponding confounds file. All resamplings can be performed with *a single interpolation step* by composing all the pertinent transformations (i.e. head-motion transform matrices, susceptibility distortion correction when available, and co-registrations to anatomical and output spaces). Gridded (volumetric) resamplings were performed using antsApplyTransforms (ANTs). Non-gridded (surface) resamplings were performed using mri_vol2surf (FreeSurfer).

**Table 1**: *Statistics and peaks for whole-brain analysis for each encoding model during sentence production. Voxelwise p < 0.005, p < 0.05 FDR cluster corrected.*

| Contrast | Cluster | | | Peak Voxel (MNI Coordinates) | | | Anatomical Location |
|---|---|---|---|---|---|---|---|
| | ***p* (FWE-corrected)** | **Size** | **mean *z*** | **x** | **y** | **z** | |
| **bag-of-nouns** | 2.60E-05 | 6217 | 0.001391 | -49 | 2 | 41 | L Precentral Gyrus |
| | 0.000286 | 1188 | 0.001447 | 2 | 12 | 51 | Supplementary Motor Area |
| | 0.011216 | 427 | 0.000847 | 26 | -68 | -51 | R Cerebellum (VIIb) |
| | 0.017631 | 330 | 0.000673 | -22 | 10 | -8 | L Putamen |
| | 0.017631 | 307 | 0.000793 | 68 | -25 | -6 | R MTG |
| | 0.035058 | 223 | 0.001185 | 61 | -30 | 29 | R Supramarginal Gyrus |
| **broad roles** | 2.24E-05 | 1984 | 0.002009 | -61 | -45 | 24 | L Supramarginal Gyrus |
| | 4.48E-05 | 1259 | 0.002311 | -51 | 2 | 41 | L Precentral Gyrus |
| | 0.002552 | 451 | 0.002305 | -36 | -60 | 56 | L Superior Parietal Lobule |
| | 0.002552 | 416 | 0.001809 | 56 | -15 | 11 | R Planum Temporale |
| | 0.002552 | 414 | 0.002134 | -2 | 12 | 54 | Supplementary Motor Area |
| | 0.021229 | 194 | 0.001937 | 1 | 32 | 36 | Posterior Cingulate Cortex |
| | 0.036216 | 147 | 0.002132 | 46 | 20 | 36 | R Middle Frontal Gyrus |
| | 0.036216 | 143 | 0.00152 | 6 | 45 | -6 | R Paracingulate Gyrus |
| | 0.037371 | 135 | 0.00234 | 41 | -72 | 36 | R Superior Parietal Lobule |
| | 0.038549 | 128 | 0.001733 | -24 | 17 | 51 | L Superior Frontal Sulcus |
| | 0.03869 | 123 | 0.001432 | 46 | -5 | -1 | R Opercular Cortex |
| | 0.040269 | 117 | 0.001267 | 26 | 67 | -51 | R Cerebellum (VIIb) |
| **narrow roles** | 0.000528 | 950 | 0.003602 | -59 | -17 | 6 | L Planum Temporale |
| | 0.005237 | 415 | 0.003266 | 3 | 27 | 56 | Supplementary Motor Area |
| | 0.018807 | 250 | 0.002346 | 61 | -45 | -11 | R MTG |
| | 0.025291 | 204 | 0.003363 | 56 | -12 | 4 | R Planum Temporale |
| | 0.025291 | 179 | 0.003387 | -2 | 25 | 39 | L Paracingulate Gyrus |
| | 0.025291 | 177 | 0.003133 | 26 | 10 | 2 | R Putamen |
| | 0.031269 | 155 | 0.003693 | -31 | 42 | 36 | L Anterior Middle Frontal Gyrus |
| | 0.032171 | 146 | 0.003239 | -14 | -30 | 74 | L dorsal Prefrontal Gyrus |

**Table 2**: *Statistics and peaks for whole-brain analysis for each encoding model during sentence comprehension. Voxelwise p < 0.005, p < 0.05 FDR cluster corrected.*

| Contrast | Cluster | | | Peak Voxel (MNI Coordinates) | | | Anatomical Location |
|---|---|---|---|---|---|---|---|
| | ***p*** **(FWE-corrected)** | **Size** | **mean** ***z*** | **x** | **y** | **z** | |
| **bag-of-nouns** | 0.000506 | 929 | 0.001026 | -54 | -40 | 26 | L Supramarginal Gyrus |
| | 0.007716 | 342 | 0.000964 | -4 | 5 | 66 | L Supplementary Motor Cortex |
| | 0.007716 | 312 | 0.00105 | -49 | 10 | 21 | L Inferior Frontal Gyrus (opercularis) |
| | 0.02089 | 200 | 0.000764 | -46 | 29 | -3 | L Inferior Frontal Gyrus (triangularis) |
| | 0.029061 | 158 | 0.00073 | -49 | -55 | -13 | L Inferior Temporal Gyrus |
| | 0.029061 | 151 | 0.000788 | 53 | 15 | 21 | R Inferior Frontal Gyrus |
| | 0.040511 | 124 | 0.000977 | -51 | 0 | 46 | L Precentral Gyrus |
| **broad roles** | 0.001772 | 753 | 0.001721 | -49 | 32 | 16 | L Inferior Frontal Gyrus (triangularis) |
| | 0.002132 | 601 | 0.001559 | -4 | 17 | 54 | L Superior Frontal Gyrus |
| | 0.003055 | 484 | 0.001522 | -56 | -65 | 11 | L Middle Temporal Gyrus |
| | 0.028618 | 204 | 0.000939 | -46 | -50 | -16 | L Inferior Temporal Gyrus |
| | 0.028618 | 196 | 0.0018 | 43 | -70 | 36 | R Lateral Occipital Cortex |
| **narrow roles** | 0.007365 | 436 | 0.003485 | -49 | 32 | 14 | L Inferior Frontal Gyrus (triangularis) |
| | 0.019496 | 273 | 0.003841 | -46 | -70 | 36 | L Angular Gyrus |
| | 0.044796 | 175 | 0.003884 | -59 | -55 | 31 | L Supramarginal Gyrus |
| | 0.044796 | 163 | 0.003748 | -11 | -75 | 49 | L Lateral Occipital Cortex |
| | 0.044796 | 151 | 0.003402 | 51 | 25 | 29 | R Middle Frontal Gyrus |

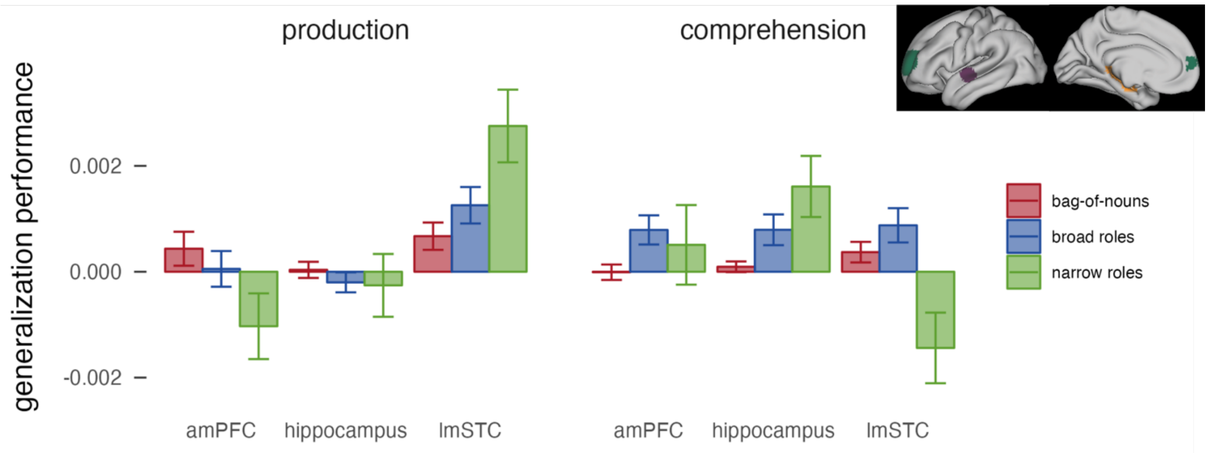

**Supplementary Figure 1**: Average generalization performance in three ROIs based on Frankland and Greene (2020a), selected independently for each participant, shown on the right. lmSTC: purple. amPFC: blue. Hippocampus: orange.
